# Voluntary movement initiation is associated with cardiac input in Libet’s task

**DOI:** 10.1101/2025.06.06.658322

**Authors:** Ksenia Germanova, Alina Studenova, Dimitri Bredikhin, Magdalena Gippert, Nikolai Kapralov, Vasily Klucharev, Arno Villringer, María Herrojo Ruiz, Vadim Nikulin

## Abstract

The relationship between motor intention and initiation of voluntary movement remains a fundamental topic in neuroscience, originating from the B. Libet seminal framework introduced in 1983. Libet’s paradigm significantly influenced discussions on intentionality, motor control, and free will. However, methodological critiques continue to challenge its interpretations, specifically the accuracy and validity of the ‘urge to move’ phenomenon. One understudied factor in this debate is the potential influence of interoceptive signals—particularly cardiac activity—in shaping the experience of motor intention and movement initiation.

In our study, we addressed this gap by examining whether cardiac signals modulate participants’ experience of the ‘urge to move’, using behavioural and electrophysiological measures in 34 healthy human participants performing Libet’s task. Crucially, when participants were asked to report the perceived ‘urge to move’, their button press timings were predominantly aligned with the diastolic phase of the cardiac cycle, indicating cardiac modulation of motor intention perception. However, analysing heart evoked potential (HEP) amplitudes as a measure of cardiac input perception, we observed no differences in HEP amplitudes associated with changes in introspective demands during the task in both source and sensor spaces.

Our results suggest that implicit perception of cardiac signals biases subjective experience of voluntary action initiation, independent from cortical interoceptive markers. These findings have implications for models of motor preparation, intentionality and the bodily basis of voluntary action, challenging conventional interpretations of motor intention and informing debates on volition and interoception.

**Significance Statement:** Our study provides evidence that implicit perception of cardiac signals influences the subjective experience of motor intention—the ‘urge to move’ in Libet’s experiment. We demonstrate, for the first time, that when reporting ‘urge to move’, participants tended to initiate voluntary movements during the diastolic phase of the cardiac cycle. These findings challenge traditional views on factors affecting motor initiation, suggesting relevance of interoceptive processing. By highlighting the role of cardiac input in experiencing motor intention, our findings impact existing debates on volition, agency and free will, further underscoring the importance of integrating bodily signals into these theoretical frameworks.

## Introduction

Understanding preparation and initiation of voluntary movements remains a fundamental challenge in cognitive neuroscience, closely tied to broader questions related to volition, intentionality, and the free will debate. At the core of this debate is Libet’s classic experiment (Libet et al., 1983), which links preparatory neural activity, the readiness potential (RP), to participants’ subjective experiences of deciding to move. Despite its significant impact, the methodology underlying Libet’s approach has been heavily criticised (Bredikhin et al., 2023; Dominik et al., 2017; Haggard & Eimer, 1999; Sanford et al., 2020). One line of critique undermines the assumption that the RP reflects a decisive aspect of movement initiation by proposing alternative interpretations of its role in motor function, decision-making and attention (for a review see Schurger et al., 2021).

The critique we focus on in this paper concerns the validity of the behavioural reports in Libet’s task (Bredikhin et al., 2023; Dominik et al., 2017; Sanford et al., 2020). Participants provide two types of timing reports after pressing a button in Libet’s task: a ‘button press’ report, marking when the movement occurred, and an ‘urge to move’ report, indicating the conscious awareness of the intention to move. Whereas button press reports rely on clear sensory feedback, the ‘urge to move’ reports are ambiguous. Previously, their accuracy and overall validity have been questioned (Banks & Isham, 2009; Bredikhin et al., 2023; Dominik et al., 2017). If present, the ‘urge to move’ may be a vague sensation that must be distinguished from ongoing sensory inputs including continuous signals from the visceral organs like the heart.

The cardiac cycle was shown to modulate visual, auditory, somatosensory, and pain perception in humans (Al et al., 2020; Duschek et al., 2013; Dworkin et al., 1994; Edwards et al., 2009; Wilkinson et al., 2013). A substantial body of evidence links the baroreceptors activation to inhibiting central processing and documents its behavioural effects (Duschek et al., 2013; Dworkin et al., 1994). Studies using carotid baroreceptor stimulation and cardiac phase analysis show improved signal detection and faster reaction times with reduced baroreceptor activation (Duschek et al., 2009; Edwards et al., 2007; Nyklíček et al., 2005). Many studies investigate central cardiac effects through heart-evoked potentials (HEPs), proposing them as a marker of cardiac interoception (Al et al., 2020; Coll et al., 2021; Kumral et al., 2022). Cardiac modulation also extends to motor functions, with evidence in health (Lai et al., 2024; Lehmann et al., 2022; Scharhag et al., 2013), and disease (Arakaki et al., 2023; Z. Wang et al., 2022). Still, the relationship between cardiac input and motor activity is not fully understood.

Previous studies examined timing of movement initiation with respect to the cardiac cycle, some suggesting facilitation during the diastole (Helin et al., 1987), and others during systole (Konttinen et al., 2003; Kunzendorf et al., 2019; Ohl et al., 2016). Findings on motor cortex excitability are also mixed: while some report no cardiac phase effects on M1 excitability (Bianchini et al., 2021; Otsuru et al., 2020), others show increased cortico-spinal excitability during systole (Al et al., 2023). The interplay between cardiac input and motor control is particularly relevant to Libet’s introspective task (Dominik et al., 2024) yet remains understudied.

In this study, we address this critical gap by examining whether cardiac input systematically biases participants’ reports and button presses in Libet’s task. Given the introspective demands of registering the ‘urge to move’, we hypothesize that the influence of cardiac input—reflected in the coupling of experiencing the ‘urge to move’ to the cardiac cycle and enhanced heart-evoked potentials (HEPs)—will be more pronounced when participants report the ‘urge to move’ compared to when they report a ‘button press’. Clarifying the interplay between cardiac signalling and introspective reports and associated button presses will refine our understanding of the factors influencing voluntary movement initiation, enhancing interpretations of neural markers like the RP and the HEP, and informing theoretical perspectives on motor control.

## Methods

### Participants

We reanalysed data from a previous study with an initial dataset containing 49 participants (Bredikhin et al., 2023). We removed 11 participants due to technical problems resulting in missing behavioural trigger information, four participants due to excessive noise in EEG. 34 remaining healthy volunteers (19 females; age range 18-57; mean age 26.27 +- 8.90) were included in this analysis. Participants were recruited through a local mailing list being either university students or regular participants of the experiments in the research centre. Participation in the experiment was compensated. All participants were naive to Libet’s clock paradigm and used their native language. The experiment was conducted following the ethical standards stated in the 1964 Declaration of Helsinki. The experimental protocol was approved by the local Ethics Committee of the HSE University (No. 19-3 dated 09 December 2019).

### Experimental Design

We used an experimental design following a modification introduced by Dominik et al. (2017) to the original Libet’s paradigm (Libet et al., 1982, 1983) (Figure 1A).

**Figure 1.**
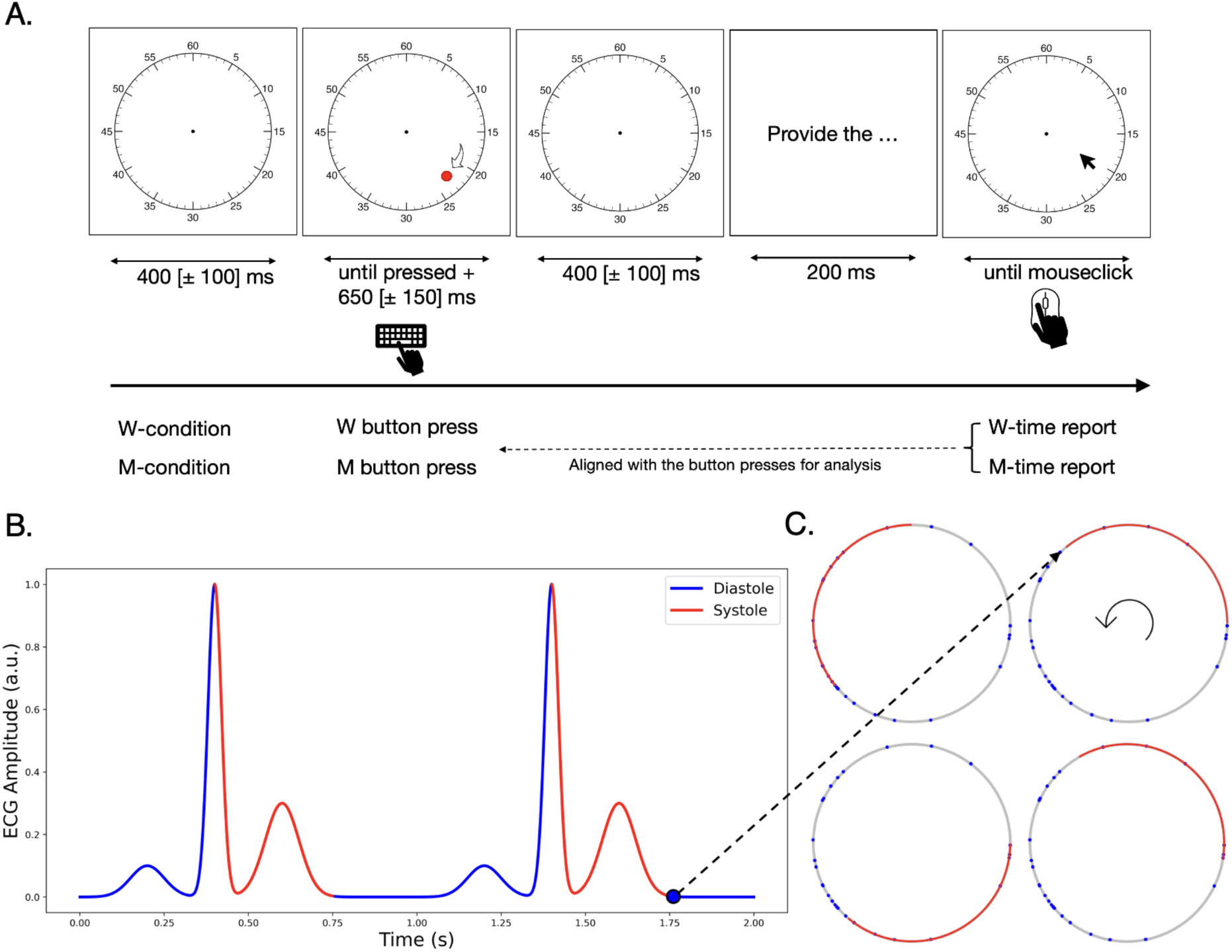
Methods. **A.** The trial structure. Each trial started with the presentation of an empty clock face, accompanied by a beep sound. After 400 [± 100] ms a red dot appeared on the clock face and started its clockwise rotation. The dot stopped when the participant pressed a space button after a random delay of 650 [± 150] ms. In both the W and M conditions, participants were instructed to spontaneously press the button focusing their gaze on the centre of the clock face. In both conditions, participants were asked to refrain from pre planning their movements. In the W condition, after a button press (W button press), participants were asked to report the time when they first experienced the intention to move (W-time report), using a mouse click. For the M-condition, after pressing the button (M button press), they had to indicate, via mouse click, the time of movement initiation (M-time report). Before the next trial, participants were instructed to rest as long as they needed. **B.** Schematic diagram illustrating the alignment process of behavioural and cardiac cycle data. To align the behavioural report of W/M button presses and W-/M-time reports (example report shown as a blue dot) with the cardiac cycle, we first determined the corresponding R–R interval. We then converted the timestamp value into a radian value, following the formula provided in the methods (see the ‘Behavioural and physiological data alignment’ section). The red colour indicates the systolic phase (R–T-offset), while the blue colour indicates the diastolic phase (T-offset–R) of the cardiac cycle. An example of how the behavioural report timestamp value is converted into radian value is shown by the dotted black arrow connecting subplots B and C. **C.** Illustration of the sliding segment test. The circle is divided into 36 equal overlapping bins, each representing a step of the systolic-like segment (shown in red) as it progresses along the circle. At each of these 36 steps, the number of observed data points (button presses in the W-condition) falling within the current segment position is calculated. This method allows for a systematic analysis of the distribution of data points relative to the segment’s length and position.

The participants were presented with two types of tasks, constituting two conditions: the W-task in the W-condition and the M-task in the M-condition. Before each condition, participants received instructions, completed three training trials, and had an opportunity to ask questions about the experimental procedure. For each trial participants were presented with a clock face. Next, a red dot appeared at a random position on a clock. A beeping sound indicated the start of the dot’s clockwise rotation. Participants were instructed to fixate their gaze at the centre of the clock to avoid following the moving dot. Participants pressed the space bar at a self-chosen moment. The dot continued rotating for 500–800 milliseconds after the keypress before disappearing. In both conditions participants were instructed not to pre-plan their movements and to act spontaneously. When the movement was completed and the dot disappeared, participants were instructed to provide a behavioural report. In the W-condition, participants were asked to report the timing of experiencing an urge to perform a movement (button press). In the M-condition, they reported the time of movement initiation. To provide these reports, participants were shown the clock face again and indicated the relevant time via a mouse click. The order of conditions was counterbalanced across participants.

Importantly, participants were not aware of the experimental instructions for the second condition while completing the first one, which made them naive to the second condition during the first part of the experiment. All participants completed 40 trials of each condition, 80 experimental trials in total. Since all movements were self-paced, participants could control the duration of the pause after each trial. The participants rested for three minutes between conditions. For further details on the experimental procedure, see Bredikhin et al. (2023).

### EEG, EMG, and ECG Recording

All signals were recorded using a 32-channel (10-20 system) EEG system (BrainAmp DC amplifier, BrainProducts). All electrode impedances were maintained below 10 kΩ. During the recording, the data were band-pass filtered between 0.1 and 100 Hz, filtered with the notch-filter for 50 Hz and sampled at 500 Hz. The recording and online pre-processing were performed using PyCoder software (BrainProducts). The eye movements (EOG) were monitored by electrodes below and from the outer canthi of the left eye. Reference electrodes were linked to mastoid processes (TP9, TP10), whereas the forehead electrode was used as the ground electrode (GND). EMG recording electrodes were placed on the first dorsal interosseous muscle of the right hand and its tendon, and the ground electrode was placed on the ulnar styloid process. The ECG was recorded using the two external channels with a bipolar ECG lead I configuration (on the forearms). All peripheral electrodes were placed after the skin was prepared with abrasive paste and alcohol wipes.

### ECG Data Processing

The ECG data were processed using Python-based custom scripts, the MNE Python toolbox (Gramfort et al., 2013, 2014) and the NeuroKit2 toolbox (Makowski et al., 2021). We divided the cardiac cycle into the systolic and the diastolic phases, defining the systolic phase as the interval from R-peak to T-wave offset, and the diastolic phase as the remaining part of the cycle (Bury et al., 2019). For the individual assessment of the onsets of the cardiac cycle phases, we determined the timings of R-peaks and T-wave offsets within the PQRST complex using algorithms from MNE Python (**mne.preprocessing.find_ecg_events**) and NeuroKit2 (**ecg_process**), respectively. The ECG data were visually inspected to ensure the correctness of parameter detection. *Behavioural and Physiological Data Alignment.* As the next step, we examined the relationship between the W- and M-time reports and button press timings with regard to the cardiac cycle. First, for both conditions we extracted timestamps of button presses and reports, aligning the latter to the movement onsets (Bredikhin et al., 2023; Libet et al., 1983). Next, we transformed all timestamps into radians in relation to R-R intervals for alignment with the cardiac cycle (Figure 1B). This transformation was achieved using the following formula:

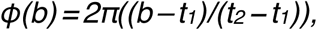

where *t_1_* and *t_2_* denote the timestamps of the R-peaks occurring before and after the behavioural timestamp *b* that we wanted to align. Consequently, *t_2_*–*t_1_* represented an R–R interval for the cardiac cycle (Helin et al., 1987; Park et al., 2020). To indicate the onset of the diastolic phase of the cardiac cycle, we individually computed the T-wave offsets for every individual and then used the mean circular value on the subject and group levels. This approach preserved the inherent ratio of systole/diastole for the cardiac cycle during the coupling process.

### EEG Data Processing

The EEG data were processed and analysed using custom Python scripts and the MNE-Python toolbox (version 1.8; Larson et al., 2024). The EEG data were high-pass filtered at 0.5 Hz and low-pass filtered at 40 Hz, then the data were visually inspected for the removal of large artefacts (e.g., movements). To remove eye movement and cardiac-field artefacts (CFA), we used independent component analysis (ICA), carried out through an FastICA algorithm (a threshold for selecting a number of principal components was 0.99, as a contrast function the logcosh was used). To ensure the correctness of ICA components selection, we used the MNE-Python toolbox functions **find_bads_ecg** and **find_bads_eog** with the z-scoring method and selected the candidate components for rejection (Dammers et al., 2008). Next, for detected components, we analysed the power spectral density (PSD) and signal coherence with the ECG and the EOG signals to ensure that only components with maximum signal coherence and similar spectral profiles were excluded from the EEG data. We removed one CFA and two ocular components for each participant.

For analysis of HEPs, the filtered signals were segmented into epochs centred around the R-peaks of the ECG signal. HEP-epochs were extracted between 200 ms prior to R-peak and 600 ms post-R-peak (Coll et al., 2021; Petzschner et al., 2019). The HEP epochs were baseline-corrected using a 100 ms interval from −200 to −100 ms of the epoch. Then the EEG epochs were inspected visually, the ones still containing excessive noise were rejected from further analysis. For between-condition comparisons, we first selected all available R-peak-related epochs from the trial onset until the button presses, yielding an average of 335 epochs for the W-condition and 309 for the M-condition available for analysis. This approach has two potential shortcomings: (1) the number of epochs differ between trials, (2) the level of engagement in interoceptive processing might also change during the trial. To account for that, we repeated the between-condition comparison including a different number of epochs per each trial, ranging from 1 epoch (last heartbeat before the button press) to the maximum number of epochs available before the button press (7 epochs).

### Statistical Analysis

To robustly analyse the distribution of latencies of behavioural responses along the cardiac cycle, we employed a Rayleigh test for uniformity at the individual-subject level, using the R **circular** toolbox (Agostinelli and Lund, 2023). For each participant, the test yielded the mean resultant length of the vector representing the distribution of latencies in each cycle, along with a *p-value* to determine whether this circular distribution deviated from uniformity. The single-subject analysis was complemented by a second-level group statistical analysis. Each participant’s distribution was represented by the mode angle on the unit circle (Galvez-Pol et al., 2022), and a group-level Rayleigh test was conducted (Al et al., 2020; Galvez-Pol et al., 2022; Park et al., 2020). We complemented this analysis with Bayesian testing using the **BayesCircIsotropy** R toolbox. We adopted the conjugate prior for the von Mises distribution, using **computeIsotropyBFHaVMConj** function with the following parameter values: R_0_ = 0 (prior resultant vector length), c_0_ = 10 (prior concentration parameter, reflecting moderate informativeness of the prior), μ_0_ (prior mean direction) was set to the mean angle of the provided sample (Mulder & Klugkist, 2021). To interpret Bayes Factors (H_1_/H_0_ as BF_10_) we referred to Wagenmakers and Wetzels (2012).

As a control analysis, we assessed the likelihood that the Rayleigh test results could be explained by a random distribution of latencies along the unit circle in our participants. We created a null distribution for each participant. For this purpose, we randomly generated angles between 0 and 2π for each trial and participant, effectively simulating M-time reports (similarly for W-time reports) that are uniformly distributed along the unit circle. We then computed the mean resultant vector for each participant based on these randomly generated angles. Following this, we combined these subject-specific mean resultant vectors to compute a group-level mean resultant vector, reflecting the collective orientation of the responses within a hypothetical uniform distribution. The Rayleigh test was then applied to this second-level data. This entire process was repeated 1000 times to construct a distribution of test statistics under the null hypothesis of uniformity. By comparing the original Rayleigh test result with this null distribution, we quantified the likelihood of observing our data assuming a uniform distribution of response latencies. We assessed whether our original Rayleigh test statistic result was larger than the values in the null distribution, at least in 95% of the cases (0.05 alpha level, one-sided test). Crucially, this analysis aims at detecting deviations from uniformity in our circular data and is independent of the systole and diastole phase length.

Next, in a second control analysis, we developed a sliding segment test to account for the contribution of the systolic and diastolic phases of the cardiac cycle into the observed non uniform data points distribution for button presses in the W-condition (Figure 1C). To achieve this, we divided the unit circle into 36 overlapping bins, each representing a potential position of the systolic-like segment. Specifically, the systolic-like segment was slid along the unit circle in incremental steps of 2π/36 radians, thereby generating 36 overlapping positions. For each segment position, we calculated the number of data points that fell within it. To evaluate whether the observed number of data points within each overlapping segment significantly deviated from chance level, we compared the observed counts per segment to expected counts given a random distribution of data points around the cardiac cycle. We established expected counts for each position by treating the probability of a datapoint falling into the systolic-like segment as proportional to the relative length of the segment (segment length divided by total circle length, 2π). Using this probability, binomial distribution parameters were defined, and critical cutoff values for statistical significance (α = 0.05) were calculated to identify segment positions with significantly higher observed counts when compared to expected counts. Additionally, we estimated the proportion of the systolic phase relative to diastole for each segment position by calculating the overlap between each shifted segment and the fixed systolic interval (from R-peak to T-wave offset). This overlap proportion was computed as the length of overlap divided by the total systolic segment length, providing a measure of systolic phase length contribution for each segment position.

To perform a within-subject comparison of HEP amplitudes between conditions in the sensor space, the HEP epochs were separately averaged for W- and M-conditions. We then performed a dependent-sample two-sided spatio-temporal cluster-based permutation test (full-epoch length, N = 1000 permutations, *alpha-level = 0.05* for initial t-test thresholding, and 0.025 for cluster selection in a two-sided test) to identify clusters with significant amplitude differences between the W- and M-condition using the MNE-Python toolbox. The same statistical procedure was applied to the source-reconstructed HEP epochs difference (see Supplemental Materials, Text S1 and Figure S1).

## Results

### Behavioural data coupling with the cardiac cycle

First, we examined the relationship between participants’ behavioural reports, button presses, and the cardiac cycle in two conditions (M and W). In the W-condition, participants were instructed to press upon feeling the intention to move (W button press) and later reported this moment (W-time report) after the button press using a mouse click. In the M-condition, they pressed at a random time of their choice (M button press) and retrospectively reported the moment of movement initiation (M-time report) (Figure 1A). Analysing data from 34 participants for the first level statistics, we observed significant Rayleigh test results in 5 participants (W-condition button presses), 3 (W-time reports), 3 (M-time reports), and 4 (M-condition button presses). The group-level statistics revealed that the distribution of button presses in the W-condition displayed a significant non-uniform distribution across the cardiac cycle (Rayleigh test *p-value* = 0.015). The direction of the group-level mean resultant vector for the button presses in the W-condition indicated an alignment to the diastolic phase of the cardiac cycle—its mean length being 0.35 (Figure 2A). The comparison to the null distribution with proportional statistics (*p-value* = 0.035) confirmed that the obtained results were not explained by a uniform distribution of events (Figure 2E). The BF analysis revealed a BF_10_ = 4.5, which corresponds to substantial evidence in favour of the alternative hypothesis H_1_. For other behavioural events, group-level analysis did not indicate a non-uniform distribution across the cardiac cycle: button presses in the M-condition (Rayleigh test, *p-value* = 0.337), W-time reports (*p-value* = 0.586), and M-time reports (*p-value* = 0.389) all showed no significant deviation from uniformity (Figure 2B-D). BF analysis (BF_10_ in 0.58–0.75 range) provided anecdotal support for the null hypothesis.

**Figure 2.**
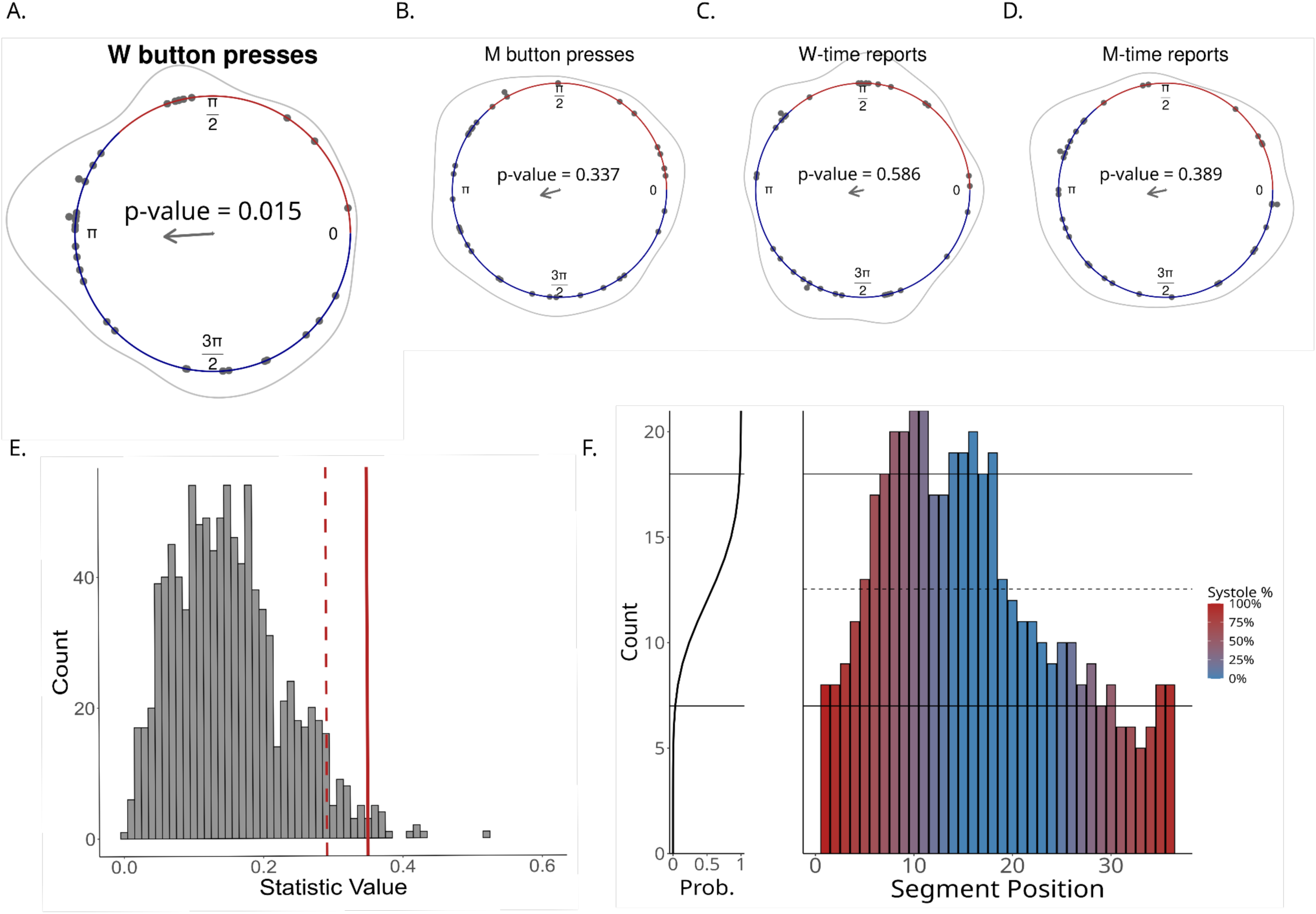
Distribution of within-subject mode W-time reports, M-time reports and button press values (n=34) across phases of the cardiac cycle. The red-coloured section of the outer circle denotes the systolic phase, while the blue-coloured line marks the diastolic phase of the cardiac cycle. The grey curve reflects the distribution of data points across the cardiac cycle; the grey dots represent within participant mode values. **A**. Button presses in the W-condition. The analysis displayed a significant non-uniform distribution across the cardiac cycle (Rayleigh test *p-value* = 0.015) with the resultant vector length being 0.35. **B.** Button presses in the M-condition. **C.** W-time reports. **D.** M-time reports. **Control analysis for behavioural data and cardiac cycle coupling**. **E**. Distribution of surrogate data and original data (red line) statistical values obtained with Rayleigh uniformity test for button presses in W-condition aligned with the cardiac cycle. The dotted red line represents the 95% distribution cutoff. The proportional statistics *p-value* = 0.035. **F.** Distribution of observed data points per segment positions assessed by the sliding segment test with horizontal dashed line representing expected mean value from the binomial distribution and solid black lines representing high and low cutoff values. The colour gradient represents the proportion of systolic (red) and diastolic (blue) phases per each segment position. The cumulative distribution plot (CDF) of the binomial distribution contextualises the observed counts relative to the expected probabilities.

Next, we evaluated the contribution of systolic and diastolic phases of the cardiac cycle to segment positions with significantly higher button-press counts than expected under a binomial distribution. We found significantly higher counts in 8 out of 36 segment positions (p-value < 0.05) (see Figure 2F). Those segment positions were predominantly located within the diastolic phase of the cardiac cycle. Overall, the evidence suggests that the non-uniform distribution of button presses in the W-condition with respect to the cardiac cycle is not due to randomness, and it highlights the predominant contribution of the diastolic phase into observed nonuniformity.

### HEP comparison between experimental conditions

Within-subject statistical analysis of differences between the W- and M-conditions in HEP amplitude including all epochs did not show any significant clusters (minimum *p-value* = 0.3) (see Figure 3). To address potential fluctuations in interoceptive processing between the trial onset and the button press, we repeated the analysis including a fixed number of epochs for each trial (ranging from maximum to minimum number of epochs available). This alternative approach yielded the same results, with no significant between-condition differences observed.

**Figure 3.**
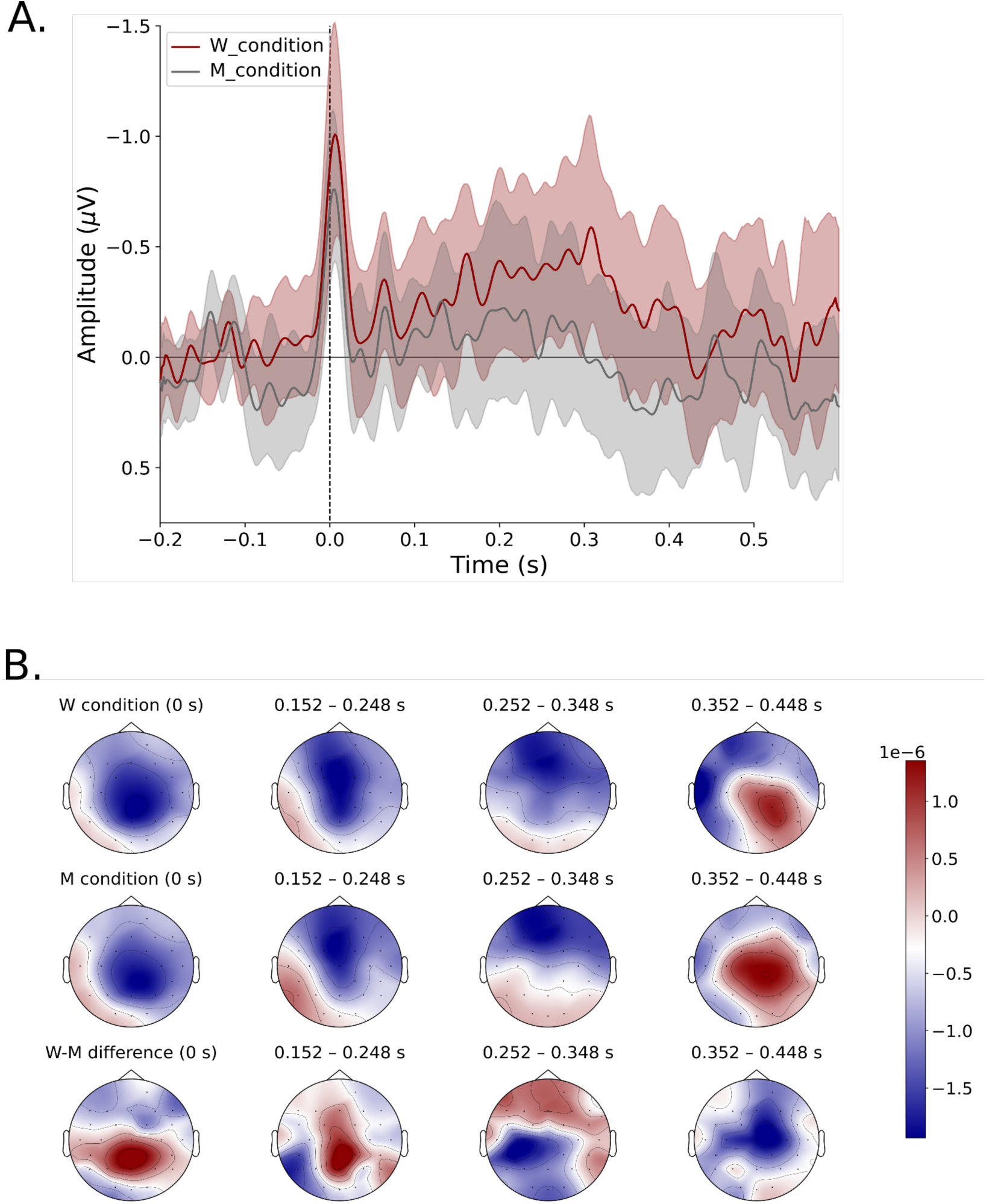
HEP amplitude in W-condition and M-condition. **A.** Descriptive ERP plot showing averages HEP for both conditions. The R-peak corresponds to 0.0 s, red colour represents the W-condition HEP, the grey–M-condition HEP, shaded area represents SEM. **B.** Descriptive topographical plots (n=34, average number of epochs in W-condition 335, in M-condition 309) for W-condition, M-condition and between-conditions difference (W *minus* M).

## Discussion

In this study, we investigated the functional role of cardiac input for movement initiation in Libet’s paradigm utilising EEG and ECG recordings to elucidate the link between visceral input and participants’ reports and button presses in Libet’s task.

We observed a non-uniform distribution for button press timings exclusively in the W-condition of the task where participants were required to report the time when they experienced the “urge to move”. W-condition button presses were predominantly aligned with the diastolic phase of the cardiac cycle. This pattern suggests a potential association between the cardiac input and participants’ detection of the ‘urge to move’, which preceded voluntary movements.

One plausible interpretation of our results is that, in the W-condition, participants exhibit increased interoceptive processing, enabling them to perceive a distinct ‘urge to move’ sensation—potentially due to a lower perceptual threshold during diastole (Al et al., 2020; Azzalini et al., 2019; Wilkinson et al., 2013). This interpretation implies that the ‘urge to move’ can be conceptualised as an internal interoceptive event accessible to conscious perception. Consequently, if the ‘urge to move’ is indeed a distinguishable internal phenomenon, its detection would require a heightened gating of interoceptive signals reflected e.g. in increased HEP amplitude. Providing that we did not observe a difference in HEP amplitude between conditions and considering that exceptionally heightened interoceptive gating would likely be necessary for detecting a subtle internal event like the ‘urge to move,’ it appears unlikely that our participants were engaging in the level of interoception required to consciously detect such sensation.

Alternatively, our findings might be explained by participants misinterpreting their cardiac signals as the ‘urge to move’. Given that this phenomenon is generally poorly defined—with no clear indication of how it should manifest—it cannot be explained to participants. Consequently, they, being unfamiliar with Libet’s task, might struggle when asked to report an ambiguous introspective event they have never consciously attended to. In such instances, they could default to a familiar, ever-present internal cue—their heartbeats—interpreting it as the ‘urge to move’.

One recent study looking into movement initiation and cardiac cycle coupling did not show any preferential alignment to the cardiac cycle for tasks involving voluntary movement initiation, including a modified version of Libet’s task (see Supplementary materials in Park et al., 2020). However, in their study participants were only presented with one condition different from both classical Libet’s task conditions (Park et al., 2020). Participants were asked to press a button as fast as possible after the rotating dot appeared on the clock face rather than being guided by any introspective event. Moreover, they were not asked to report any subjective timings, which altogether makes this task less introspective than the one we implemented.

Interestingly, we did not observe any evidence of the W-time reports relatedness to the cardiac cycle. The button presses in the W-condition and the W-time reports are likely to be generated by different processes. In the W-condition, participants are required to react when detecting an internal trigger—‘urge to move’—by pressing a button while focusing their gaze on the centre of the clock face. Specifically, participants are instructed to avoid looking at the numbers on the clock face during the experimental trial to prevent anchoring their movements to these visual cues. In contrast, reporting the W-time requires reproducing the time of the ‘urge to move’ in the units of the clock (“seconds”). The complexity of reporting the W-time may lead to a significant variability between the button press and subsequently reported W-time value. This variability could account for the absence of the W-time reports alignment with the cardiac cycle. The W-time reports were previously shown to be influenced by the manipulation of Libet’s clock parameters (e.g., the rotation speed and the number of clock markings (Ivanof et al., 2022), and by the order of the task condition presentation (Bredikhin et al., 2023; Dominik et al., 2017). Taken together, these findings suggest that participants might act upon their cardiac input interpreting it as the ‘urge to move’, which manifests in locking W-condition button presses to the cardiac cycle and question the overall reliability of the W-time reports.

We did not observe a significant difference in HEP amplitudes between the conditions of the task. One explanation for the absence of HEP effects may lie in methodological considerations—specifically, the limited number of epochs and thus lower signal-to-noise ratio (SNR) for evoked responses, which may have reduced sensitivity to between-condition amplitude differences. Given the relatively small HEP amplitudes reported in previous studies and the potential fluctuations in interoceptive state throughout the task, capturing an HEP effect is challenging. In our design, increasing the number of HEP epochs requires extending the time window from which R-peaks are selected, which may include periods further away from the critical event (i.e. the button press). This introduces a trade-off: using fewer epochs improves temporal specificity but reduces SNR, while including more epochs enhances SNR at the cost of potentially diluting the effect by averaging over periods with lower interoceptive engagement.

While the HEP is frequently assumed to be an objective measure of engagement in interoceptive processing (Buot et al., 2021; Kumral et al., 2022; Poppa et al., 2022; Verdonk et al., 2024), the relationship between the HEP and interoception is not fully clarified. Reported clusters of HEP differences are heterogeneous in terms of their spatio-temporal localization with multiple confounding factors. Coll et al. (Coll et al., 2021) highlight in their systematic review that observed associations between HEP dynamics and interoceptive manipulations often suffer from considerable methodological heterogeneity. Crucially, studies differ significantly regarding their control for the pulsatility artifact, a factor potentially confounding the HEP characteristics. With proper control for the pulsatility artefact, some studies report that previously observed HEP differences vanish (Buot et al., 2021). Thus, caution is needed when interpreting HEP amplitude modulation as direct evidence of heightened interoceptive processing.

The absence of between-condition HEP differences in our study may reflect a lack of significant cardiac output modulation at the cortical level. Many studies reporting differences in HEPs utilise designs involving modulation of cardiac output. For instance, significant HEP differences are often observed when comparing healthy individuals with clinical cohorts characterized by altered cardiac parameters such as heart rate variability (HRV) or stroke volume (Kumral et al., 2022; Lutz et al., 2019; Yoris et al., 2018). Additionally, studies employing pharmacological interventions or stimulations that affect cardiac function reliably demonstrate pronounced HEP effects (Pollatos et al., 2016; Poppa et al., 2022; Schulz et al., 2015). Furthermore, tasks explicitly designed to amplify participants’ attention towards their own heart, which could also alter cardiac parameters, display HEP modulations (Coll et al., 2021). In contrast, our experimental design involves healthy individuals and a task that does not explicitly direct participants’ attention towards cardiac activity, nor does it involve cardiac parameters manipulation. Given that cardiac awareness was entirely implicit in our study, we did not anticipate substantial changes in cardiac parameters between experimental conditions.

Moreover, recent evidence suggests that the neural correlates of cardiac signals extend beyond cortical processing. Cardiac input is consistently relayed through subcortical and peripheral nervous structures, mediated primarily by Piezo-dependent mechanoreceptors located in the heart, arteries, skeletal muscle, and chest wall (Hamill, 2023; Salameh et al., 2024). These signals, transmitted via the vagus nerve, glossopharyngeal, and spinal nerves, reach brainstem and thalamic regions before ascending to the cortical areas associated with autonomic and somatosensory processing (Hamill, 2023; Park & Blanke, 2019). Notably, cardiac input is continuously present since early development, suggesting it might predominantly function through implicit modulation of neural dynamics rather than resulting in explicit conscious perception of cardiac input (Hamill, 2023). It may influence cortical neuronal dynamics through vascular-neural coupling (VNC) mechanisms, affecting neuronal activity with mechanical pulsations of intracranial vessels by means of both slow autoregulatory and rapid pressure-activated mechanisms (Hamill, 2023; J. Wang & Hamill, 2021). Salameh et al. (2024) provided evidence from animal studies supporting the neuronal origin of slow local field potential oscillations coupled to vascular pulsations. Collectively, these findings indicate that cortical cardiac effects, reflected in HEP, might not be a prerequisite for observing behavioural cardiac-related changes.

In future studies, to further disentangle the alternative explanations of our results, it would be important to separately measure cardiac interoception using a reliable heartbeat perception task and relate it to the strength of the button-press allocation to the cardiac cycle in the W-condition.

In conclusion, our study contributes to the understanding of interoceptive processing in the context of voluntary movement initiation, particularly emphasising the role of the cardiac input. The observed non-uniform distribution of button presses aligning predominantly with the diastolic phase of the cardiac cycle, highlights a connection between cardiac input and motor initiation in the W-condition. Additionally, our results further challenge previous interpretations of the W-time and support methodological and conceptual criticism of the ‘urge to move’ phenomenon. Our findings also underscore the unresolved debate regarding the reliability of HEP as a direct measure of interoceptive engagement.

## Code availability

The code is available via the open Github repository: https://github.com/KseniaGermanova/Libet-HEP-paper

## Author contribution

K.G. and D.B. collected the data; D.B., V.N. and V.K. designed the experiment; V.N., M.H.R., A.S., N.K. and M.G. conceived ideas for additional data analyses; K.G. analysed the data; M.G. and N.K. reviewed the code; K.G. prepared the original draft; K.G., M.H.R., V.N., N.K., M.G., A.V., V.K., D.B. and A.S. reviewed and edited the manuscript; M.H.R. and V.N. supervised the project.

## Acknowledgments

The authors thank the International Max Planck Research School on Cognitive NeuroImaging (IMPRS CoNI) for supporting Ksenia Germanova financially.

V.K. was partially funded by the Basic Research Program at the HSE University.

