## Supplemental Materials Text S1 and Figure S1 for "Voluntary movement initiation is associated with cardiac input in Libet’s task"

Text S1.

*Source analysis. Methods.* Source reconstruction was implemented using MNE-Python functions and pipelines (Gramfort et al., 2014). For source localisation, we used the difference between average HEPs for W- and M-condition and ‘fsaverage’ subject from FreeSurfer (Fischl, 2012). The forward model was computed using the 3-layer Boundary Element Method (BEM) with 4098 voxels per hemisphere. Next, we applied eLORETA (Pascual-Marqui et al., 2011) for source reconstruction with the following parameters: normal orientation of sources to the cortical surface, regularisation parameter set to 0.05, and identity noise covariance matrix.

*Source analysis. Results.* To determine the source localization of the HEP epochs, we performed source reconstruction analysis of the epochs in W- and M-conditions. On the reconstructed data, we implemented a similar statistical procedure (see *Methods, Statistical analysis*) to account for potential between-condition differences in reconstructed sources, however the test did not find any significant clusters (the smallest cluster *p-value* = 0.3, Figure S1).

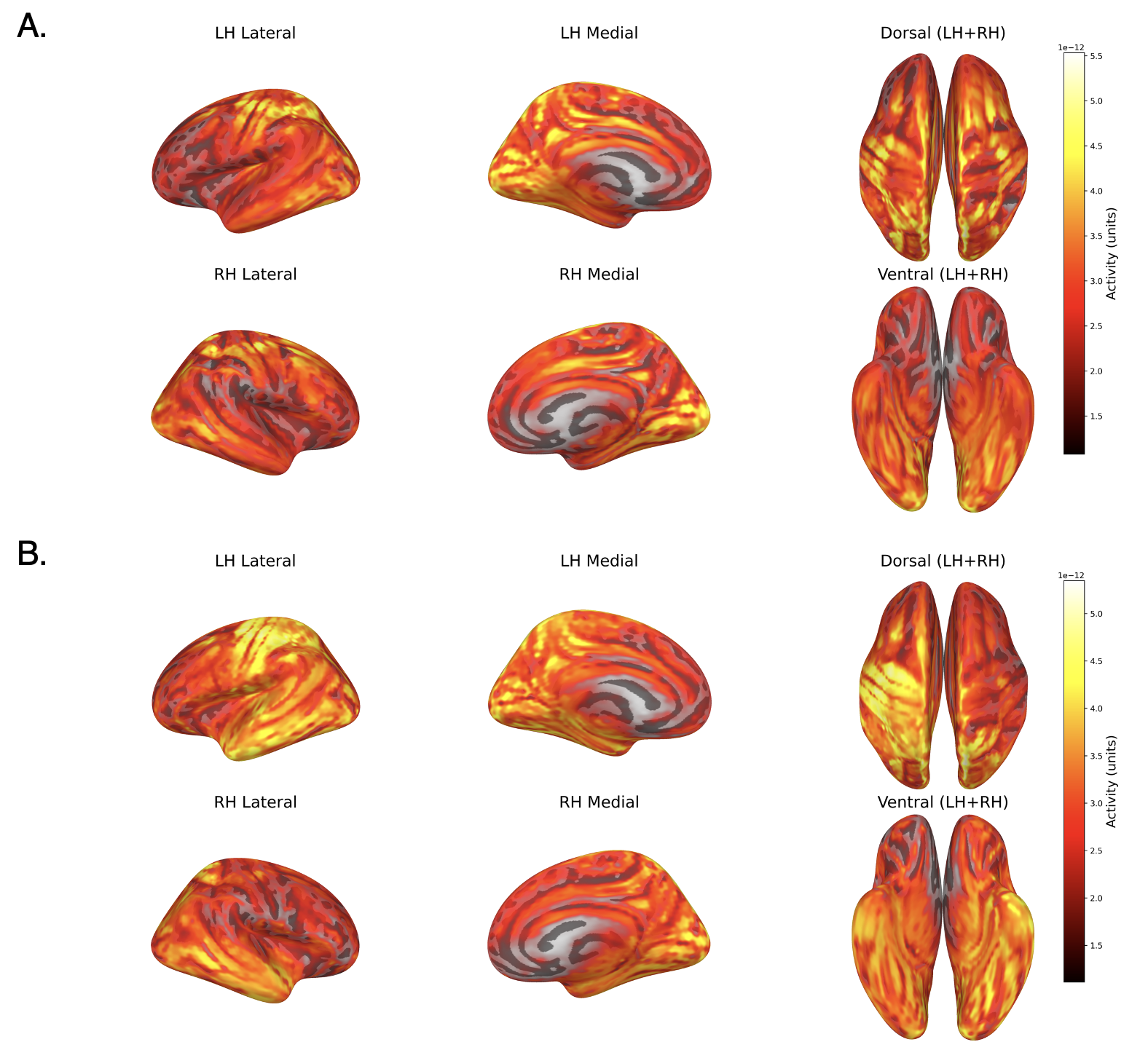

**Figure** **S1**. **HEP source reconstruction**. **A.** Source reconstructed HEP epochs in the W-condition (absolute values) at 0.2 seconds after the R-peak. **B.** Source reconstructed HEP epochs in the M-condition (absolute values) at 0.2 seconds after the R-peak.
